# Probing *E. coli* SSB Protein-DNA topology by reversing DNA backbone polarity

**DOI:** 10.1101/2020.12.05.412478

**Authors:** Alexander G. Kozlov, Timothy M. Lohman

**Author notes:** Address correspondence to: Department of Biochemistry and Molecular Biophysics, Box 8231, Washington University in St. Louis School of Medicine, 660 South Euclid Ave. St. Louis, M0 63110.

## Abstract

*E. coli* single strand (ss) DNA binding protein (SSB) is an essential protein that binds ssDNA intermediates formed during genome maintenance. SSB homo-tetramers bind ssDNA in two major modes differing in occluded site size and cooperativity. The (SSB)_35_ mode in which ssDNA wraps on average around two subunits is favored at low [NaCl] and high SSB to DNA ratios and displays high “unlimited”, nearest-neighbor cooperativity forming long protein clusters. The (SSB)_65_ mode, in which ssDNA wraps completely around four subunits of the tetramer, is favored at higher [NaCl] (> 200 mM) and displays “limited” low cooperativity. Crystal structures of *E. coli* SSB and *P. falciparum* SSB show ssDNA bound to the SSB subunits (OB-folds) with opposite polarities of the sugar phosphate backbones. To investigate whether SSB subunits show a polarity preference for binding ssDNA, we examined *Ec*SSB and *Pf*SSB binding to a series of (dT)_70_ constructs in which the backbone polarity was switched in the middle of the DNA by incorporating a reverse polarity (RP) phosphodiester linkage, either 3’-3’ or 5’-5’. We find only minor effects on the DNA binding properties for these RP constructs, although (dT)_70_ with a 3’-3’ polarity switch shows decreased affinity for *Ec*SSB in the (SSB)_65_ mode and lower cooperativity in the (SSB)_35_ mode. However, (dT)_70_ in which every phosphodiester linkage is reversed, does not form a completely wrapped (SSB)_65_ mode, but rather binds *Ec*SSB in the (SSB)_35_ mode, with little cooperativity. In contrast, *Pf*SSB, which binds ssDNA only in an (SSB)_65_ mode and with opposite backbone polarity and different topology, shows little effect of backbone polarity on its DNA binding properties. We present structural models suggesting that strict backbone polarity can be maintained for ssDNA binding to the individual OB-folds if there is a change in ssDNA wrapping topology of the RP ssDNA.

**Statement of Significance:** Single stranded (ss) DNA binding (SSB) proteins are essential for genome maintenance. Usually homo-tetrameric, bacterial SSBs bind ssDNA in multiple modes, one of which involves wrapping 65 nucleotides of ssDNA around all four subunits. Crystal structures of *E. coli* and *P. falciparum* SSB-ssDNA complexes show ssDNA bound with different backbone polarity orientations raising the question of whether these SSBs maintain strict backbone polarity in binding ssDNA. We show that both *E. coli* and *P. falciparum* SSBs can still form high affinity fully wrapped complexes with non-natural DNA containing internal reversals of the backbone polarity. These results suggest that both proteins maintain a strict backbone polarity preference, but adopt an alternate ssDNA wrapping topology.

## Introduction

Single-stranded DNA binding proteins (SSBs) are essential in all kingdoms of life and function by selectively binding to the single-stranded DNA (ssDNA) intermediates that are formed during genome maintenance protecting them from degradation and inhibiting DNA secondary structures (1–4). *Escherichia coli* SSB (*Ec*SSB) also serves as a central hub for interaction with numerous metabolic proteins (SSB interacting proteins – SIPs) involved in replication, recombination and repair (5).

*Ec*SSB functions as a homo-tetramer (Fig. 1B) (3, 6), with each subunit (177 amino acids) composed of two domains (Fig. 1A): a structured N-terminal DNA binding domain (DBD) (residues 1-112), and a C-terminal domain (residues 113-177) composed of a flexible, intrinsically disordered linker (IDL) [56 aa (Fig. 1A)] and a nine residue “acidic tip”. This acidic tip is conserved among many bacterial SSBs and is the primary site of interaction with the SIPs (5, 7–13). *Ec*SSB binds ssDNA in two major modes referred to as (SSB)_35_ and (SSB)_65_, where the subscripts denote the average number of ssDNA nucleotides occluded (14, 15). The relative stabilities of these modes depend primarily on salt concentration and type, and protein to DNA ratio (binding density) (14, 16–24), as well as applied force (25–28).

**Figure 1.**
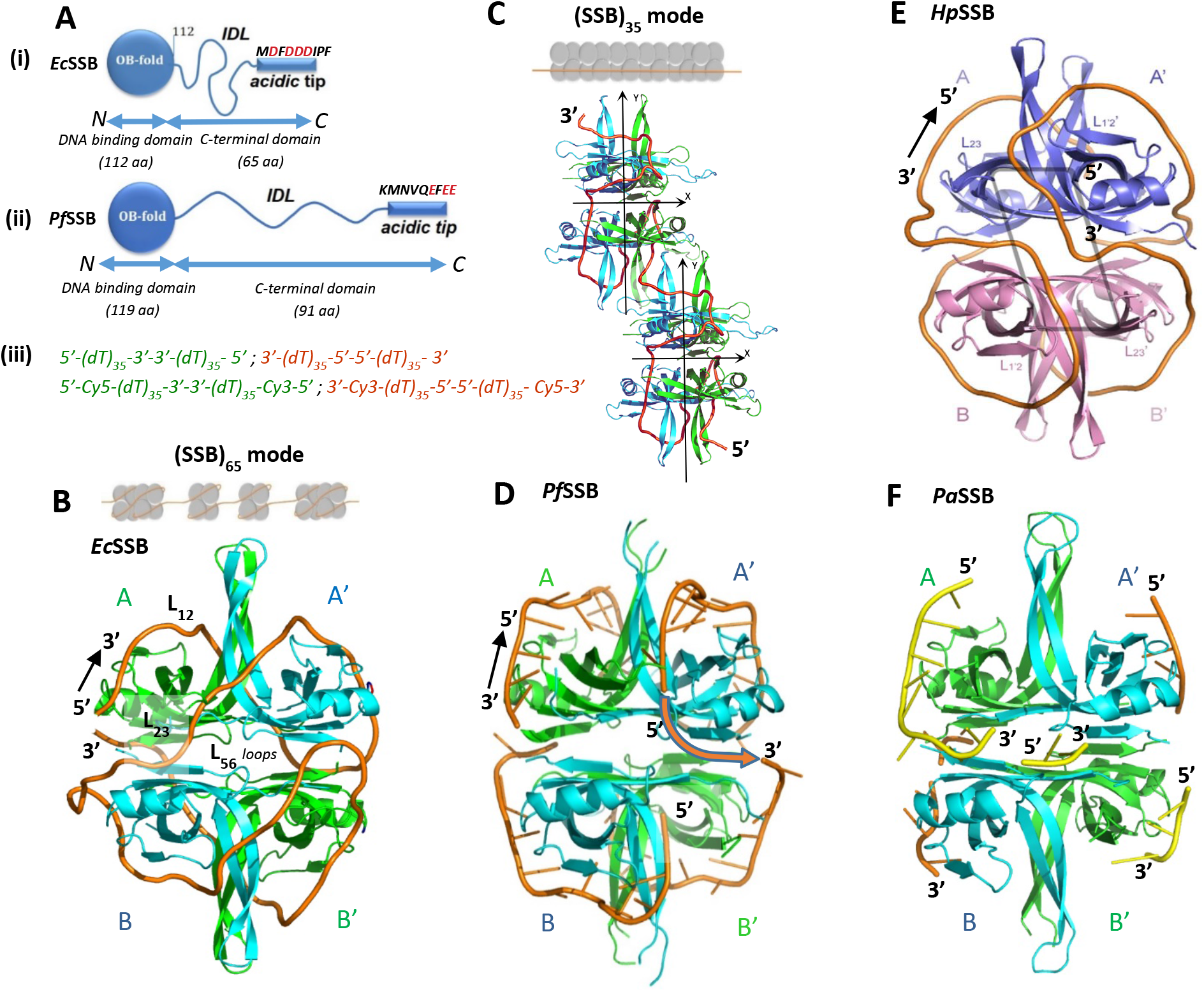
*E. coli* SSB-ssDNA structure and binding modes and its comparison with other SSB-DNA structures. **(A)** – *Ec*SSB (i) subunit (177 aa) and *Pf*SSB (ii) subunit (210 aa) are composed of an N-terminal DNA binding domain (OB fold) (112 aa and 119 aa, respectively), intrinsically disordered linkers (IDL), 56aa and 82aa long, respectively, and 9aa acidic tips, with different aa sequences. (iii) – RP (reverse polarity) (dT)_70_ constructs used in the study in which the polarity of phosphodiester backbone was reversed in the middle of the DNA (abbreviated as 5’-(dT)_70_-5’ (green) and 3’-(dT)_70_-3’ (dark orange)). **(B)** – cartoon of *Ec*SSB-ssDNA complex in its (SSB)_65_ binding mode (6), with 65 nts of DNA (orange ribbon) wrapped around the tetramer, showing the polarity of DNA wrapping. The identical subunits are marked as AA’BB’. Note that the IDL and acidic tip are not visible in this structure. **(C)** – cartoon of a proposed model for *Ec*SSB-ssDNA binding to (dT)_70_ (orange tube) in the (SSB)_35_ binding mode using an average of only two subunits per tetramer. **(D)** – cartoon of a *Pf*SSB in complex with two molecules of (dT)_35_ (orange tubes) showing a polarity for ssDNA binding that is opposite to *Ec*SSB (41). **(E)** – cartoon of *Hp*SSB-ssDNA complex in the (SSB)_65_ binding mode (42). **(F)** –1 cartoon of *Pa*SSB in complex with two molecules of (dT)_25_ (44).

In the (SSB)_65_ mode, favored at [NaCl]>0.20 M or [Mg^2+^]>10 mM at 25°C, the ssDNA wraps around all four subunits of the tetramer (6) (see Fig. 1B), with a ~65 nucleotide occluded site size. The topology of ssDNA wrapping in the (SSB)_65_ binding mode follows the seams on a baseball such that ssDNA enters and exits the tetramer in close proximity. On long ssDNA, the (SSB)_65_ mode displays “limited” cooperativity between adjacent tetramers (16, 24, 29) as depicted in Fig. 1B. In this mode SSB can diffuse along ssDNA transiently destabilizing DNA hairpins and promoting RecA filament formation (25, 30). Single stranded DNA translocases are able to actively push SSB tetramers bound in the (SSB)_65_ mode along ssDNA providing a potential mechanism for reorganization and clearance of tightly bound SSBs from ssDNA (31). In the (SSB)_65_ mode, non-nearest neighbor SSB tetramers can interact cooperatively, possibly through IDLs (23, 32). These interactions result in condensation of nucleoprotein complexes and require presence of IDL and are promoted by glutamate and acetate salts (23, 28, 32).

In the (SSB)_35_ mode, favored at [NaCl]<10 mM (Fig. 1C) or [MgCl_2_]<1 mM, and high SSB to DNA ratios (14, 15, 18), ssDNA wraps around only two subunits on average with a ~35 nucleotide occluded site size. In this mode SSB binds ssDNA with unlimited nearest-neighbor cooperativity favoring formation of long protein clusters (17, 18, 20, 22, 24, 32, 33) as depicted in Fig. 1C. Structural models for the (SSB)_35_ binding mode have been proposed, suggesting direct interactions of adjacent tetramers through the L_45_ loops within the tetrameric core of the protein (6) (Fig. 1C). In this mode SSB can diffuse along ssDNA (25, 26) and undergo direct or intersegment transfer between separate ssDNA molecules (34) or between distant sites on the same DNA molecule (35). The ability to undergo direct transfer appears to play a role in SSB recycling during replication (34, 36). Nearest neighbor cooperativity can also be displayed between the different SSB binding modes (24). A recent study (37) identified residues Y22 and K73 as important for cooperative interactions between adjacent tetramers. Both the linker and the conserved tip of C-terminus are also important for cooperativity (23, 24, 32).

Most bacterial SSBs are homo-tetramers (38). Although the dimeric *Deinococcus radiodurans* SSB (DrSSB) is an exception (39, 40), it still contains 4 OB folds (2 OB folds per monomer). In addition to *E. coli*, several other homotetrameric SSB-ssDNA crystal structures have been reported, including *Plasmodium falciparum* (*Pf*SSB) (41), *Deinococcus radiodurans* (*Dr*SSB (40), *Helicobacter Pylori* (*Hp*SSB) (42), *Bacillus subtilis* (*Bs*SSB-B (43) and *Bs*SSB-A (37)), and *Pseudomonas aeruginosa* SSB (*Pa*SSB)(44). All but one of these structures shows DNA bound to the OB-folds with the same phosphodiester backbone polarity orientation (following the path across the β-sheet in the direction from 2-3 loop towards 1-2 loop). This is exemplified by *Pf*SSB and *Hp*SSB in Fig. 1D and 1E. The one exception is the *Ec*SSB-DNA structure that displays the opposite 5’-3’ polarity to each OB-fold (Fig. 1B). It is interesting to note that the predicted topology of ssDNA wrapping in the (SSB)_65_ binding mode also differs for *Ec*SSB when compared to *Pf*SSB and *Hp*SSB. A crystal structure of *Ec*SSB tetramer bound with two (dC)_35_ molecules (6) predicts a ssDNA binding topology that wraps around the OB folds of the upper AA’ dimer (first (dC)_35_ molecule), then crosses the dimer-dimer interface in the direction of the B subunit α-helix and finally wraps around the OB folds of the BB’ dimer (second (dC)_35_ molecule) as shown in Fig. 1B (suggested path is AA’BB’). In a *Pf*SSB structure (41), the first and the second molecule of (dT)35 are wrapped around the upper and lower dimer, similarly to *Ec*SSB (though with different polarity). However, crossing of the dimer-dimer interface occurs in the direction of the B’ subunit following a path under the α-helix of the A’ subunit as shown in Fig. 1D (suggested path is AA’B’B). Essentially, the same path is predicted for the (SSB)_65_ mode of *Hp*SSB (42)(Fig. 1E) and *Bs*SSB-B (43). Interestingly, a recent structure of *Pa*SSB co-crystallized with (dT)_25_ suggests that individual subunits could accommodate ssDNA with opposite polarity (e.g. (dT)_25_ shown in yellow in Fig. 1F binds to subunit A with 5’-3’ polarity, whereas it binds to subunit B’ with 3’-5’ polarity).

Based on available SSB-DNA structural data it is reasonable to expect that polarity of ssDNA binding might affect the SSB wrapping topology and the binding properties. In the current study we examined the effects of ssDNA backbone polarity for *Ec*SSB and *Pf*SSB for two reasons. First, crystallographic studies suggest that they bind ssDNA with opposite polarities and slightly different topologies. Second, their ssDNA binding properties have been extensively studied in solution. In particular, it was established that *Pf*SSB forms a (SSB)_65_ binding mode similar to that of *Ec*SSB, yet *Pf*SSB does not form a stable (SSB)_35_ binding mode (45). Yet, both proteins can diffuse on ssDNA in their fully wrapped modes, and effectively melt out short hairpins (25, 30, 45). Here we examine the binding of these proteins to a variety of oligodeoxythymidylates (dT_70_) that contain a reversal of the phosphodiester backbone polarity in the middle of the DNA molecule (after residue 35) either through a 3’-3’ or 5’-5’ linkage. We also examine the binding of ap-(dT)_70_ and ap-(dT)_35_ DNA in which the polarity alternates after every residue.

## Materials and Methods

### Reagents and buffers

Buffers were prepared with reagent grade chemicals and distilled water treated with a Milli Q (Millipore, Bedford, MA) water purification system. Buffer T is 10 mM Tris, pH 8.1, 0.1 mM Na_3_EDTA; buffer H is 10 mM Hepes, pH 8.1, 0.1 mM Na_3_EDTA.

### SSB proteins and ssDNA

*E. coli* SSB protein (SSB) was expressed and purified as described (32). The SSB protein is a stable tetramer under all solution conditions used in this study as determined by sedimentation velocity (32). Protein concentration was determined spectrophotometrically (14) (buffer T, 0.20 M NaCl) using ε_280_=1.13 × 10^5^ M^-1^ cm^-1^ (tetramer). *Plasmodium falciparum* SSB (*Pf*SSB) was expressed and purified as described (41) and its concentration was determined using ε_280_=9.58× 10^4^ M^-1^ cm^-1^ (tetramer) (41, 45)

Oligodeoxythymidylates possessing 70 and 35 nucleotides were used in this study ((dT)_70_ and (dT)_35_, respectively). Along with DNA possessing normal 5’ to 3’ polarity, we also examined two variations of (dT)_70_ (Fig. 1A(iii)) in which the backbone polarity was reversed in the middle of the sequence: 5’-(dT)_35_-3’-3’-(dT)_35_-5’ (abbreviated as 5’-(dT)_70_-5’) and 3’-(dT)_35_-5’-5’-(dT)_35_-3’ (abbreviated as 3’-(dT)_70_-3’). Two DNA molecules, termed ap-(dT)_70_ and ap-(dT)_35_ were synthesized in which the backbone polarity is reversed after each residue. A similar set of (dT)_70_ molecules were synthesized that were doubly labeled at the ends with Cy5 and Cy3 fluorescence dyes: (5’-Cy5-(dT)_70_-Cy3-3’, 5’-Cy5-(dT)_35_-3’-3’-(dT)_35_-Cy3-5’, 3’-Cy5-(dT)_35_-5’-5’-(dT)_35_-Cy3-3’ and 5’-Cy5-ap-(dT)_70_-Cy3-3’). All oligonucleotides were synthesized and purified to >99% homogeneity as described (20, 46). DNA concentrations were determined spectrophotometrically in buffer T + 0.10 M NaCl using ε_260_ = 8.1×10^3^ M^-1^ (nucleotide) cm^-1^ for all not labeled (dT)_70_ and (dT)_35_ constructs (47) (70×ε_260_ and 35×ε_260_, molecule). For all Cy5-(dT)_70_-Cy3 the extinction coefficient ε_260_ = 5.82×10^5^ M^-1^ (molecule) was used (24, 46).

### Fluorescence equilibrium titrations

Fluorescence titrations were performed in buffers T and H, 25°C, at the NaCl and NaBr concentrations indicated in the text and Figure legends, using a QM-4 spectrofluorometer (Photon Technology International/Horiba Scientific, Edison, NJ). Reverse titrations of SSB (0.3 μM) with (dT)_70_ constructs (Fig. 2,4 and 6) and normal titration of SSB (0.3 μM) with ap-(dT)_35_ (Fig. S5) were performed by monitoring quenching of the intrinsic SSB tryptophan fluorescence (λ_exc_=296 nm and λ_em_=345 nm) and analyzed as described (24, 32, 48). Normal titrations of Cy5-(dT)_70_ Cy3 constructs (0.1 μM) with SSB (Fig. 2,3,6 and Fig. S2) were performed by exciting Cy3 donor (515 nm) while monitoring sensitized emission from Cy5 acceptor at 665 nm and analyzed as described (21, 24, 48).

**Figure 2.**
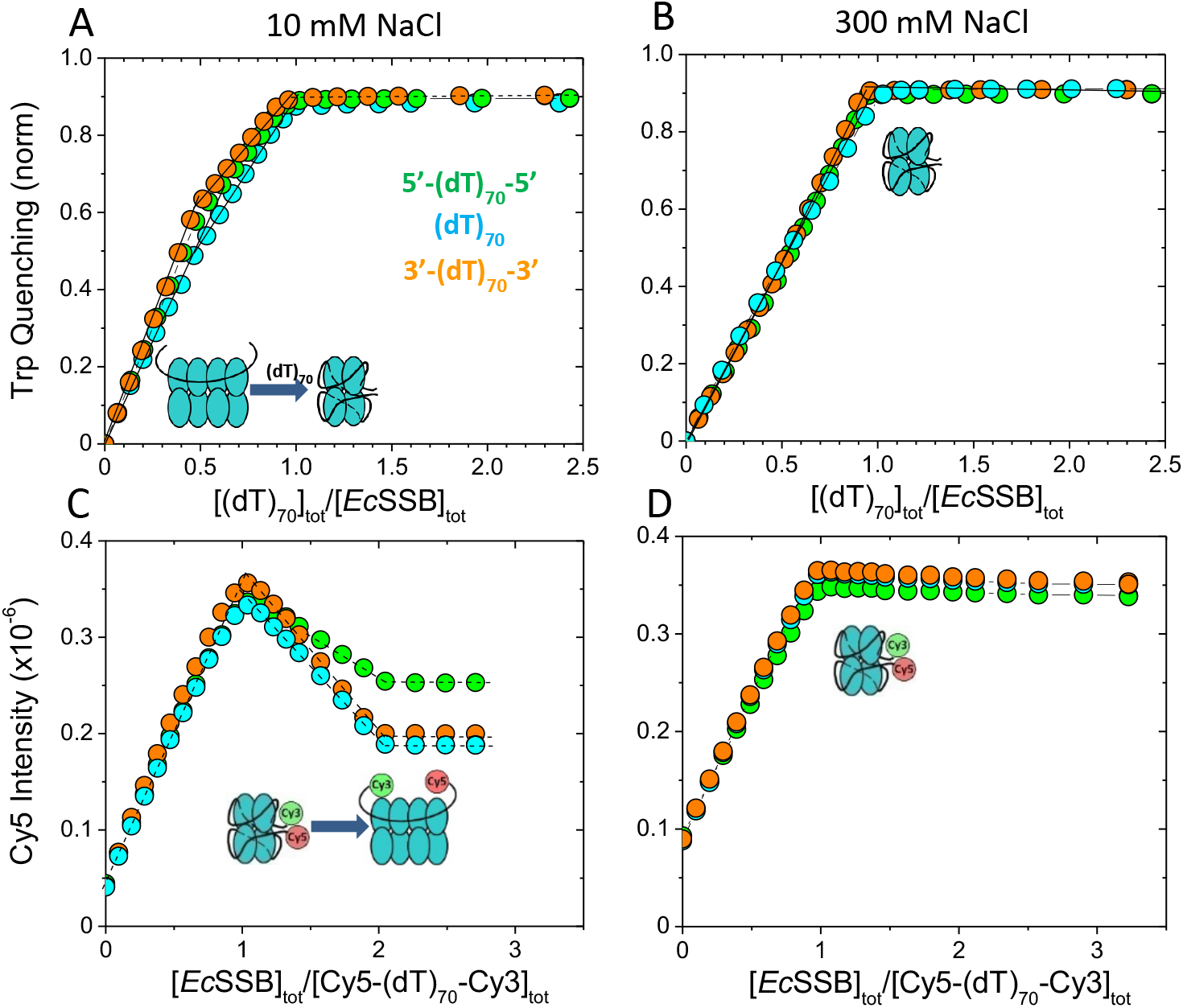
*Ec*SSB can form (SSB)_35_ and (SSB)_65_ modes on reverse polarity (dT)_70_. **(A)** and **(B)** - results of equilibrium titrations of *Ec*SSB (0.30 μM) with (dT)_70_ (cyan), 3’-(dT)_70_-3’ (orange) and 5’-(dT)_70_-5’ (green) (ex: 296 nm, em: 350 nm) in 10 mM NaCl and 0.30 M NaCl (buffer-T, 25°C), plotted as fraction of Trp fluorescence quenching versus the ratio of total DNA to total SSB tetramer concentrations. The cartoons depict the (SSB)_35_ and (SSB)_65_ binding modes that form during the course of the titrations. **(C)** and **(D)** - results of equilibrium titrations of Cy3/Cy5 labeled (dT)_70_ (cyan), 3’-(dT)_70_-3’ (orange) and 5’-(dT)_70_-5’ (green) (0.1 μM each) (see Figure 1A(iii)) with *Ec*SSB monitoring Cy5 fluorescence enhancement (exc: 515 nm, em: 665 nm) in 10 mM NaCl and 0.30 M NaCl (buffer-T, 25°C). The cartoons depict the (SSB)_35_ and (SSB)_65_ binding modes and where they form during the course of the titrations.

Binding isotherms in Fig. 4 for (dT)_70_ (X) binding to SSB (M) to determine equilibrium binding constants (*K*) were analyzed using a single site binding model described in Eq. 1 (48),

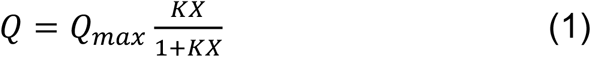

where *Q=(F_0_-F_obs_)/F_0_* is normalized Trp fluorescence quenching with *F_0_* representing the fluorescence intensity of SSB alone and *F_obs_* the Trp fluorescence measured at each point in the titration; and *Q_max_=(F_o_-F_max_)/F_o_* is the normalized fluorescent quenching at saturation. The concentration of the free DNA (*X*) in Eq. 1 was obtained by solving mass conservation Eq. (2):

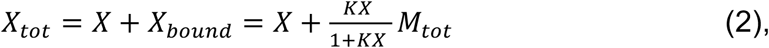

where X_tot_ and M_tot_ represent total concentrations of (dT)_70_ and SSB, respectively. Non-linear least squares (NNLS) fitting of the isotherms to Eqs. 1–2 was performed as described (48) using SCIENTIST (Micromath, St. Louis, MO)

### Isothermal titration calorimetry

ITC titrations were performed in buffer H (25°C, 20 mM NaCl) using VP-ITC microcalorimeter (Malvern Panalytical Inc., Wesborough, MA). In these experiments the SSB (1μM) was titrated with (dT)_70_ DNA constructs with stock solutions of reagents thoroughly dialyzed prior use. The data were analyzed using software provided by manufacturer as described (49)

### Stopped-Flow kinetics

Fluorescence stopped-flow experiments were performed as described (46) in Buffer H, 25°C, in the presence of 2M NaBr using an Applied Photophysics SX.18MV instrument (Applied Photophysics Ltd., Leatherhead, U.K.). In experiments shown in Fig. S4 the SSB at concentration 50 nM (final) was mixed with increasing concentrations of (dT)_70_ constructs and intrinsic T rp florescence (λ_exc_=296 nm) signal change was monitored at (λ>340 nm) using long-pass filter (Oriel, catalog no. 51258). The observed rates, r_obs_ (s^-1^), obtained fitting signal change to one-exponential decay function were than plotted as a function of DNA concentration (Fig. S4). Linear fit of these data produced kon and koff as slope and intercept, respectively.

### Analytical sedimentation

Sedimentation velocity experiments were performed as described (23, 24, 32) with an Optima XL-A analytical ultracentrifuge and An50Ti rotor (Beckman Instruments, Fullerton, CA) at 42000 rpm in buffer H, 25°C, with the salt concentrations and type indicated in the text and Fig. S1 and S3 legends. Constant concentrations of (dT)_70_ constructs (0.5 μM) were used while adding SSB to make solutions with different protein to DNA ratios (r=[SSB]_tot_/[(dT)_70_]_tot_). The absorbance was monitored at 260 nm, reflecting predominantly the contribution from ssDNA. The contribution of SSB to the absorbance at 260 nm is small compared to the DNA at protein/DNA ratios, r≤2, used in this study. Data were analyzed using SEDFIT (www.analyticalultrasentrifugation.com) to obtain c(s) distributions and estimate molecular weights for the species described by single-peaks (50). The c(s) refers to the continuous sedimentation distribution that results from a SEDFIT analysis of the absorbance traces obtained in a sedimentation velocity AUC experiment using the Lamm equation (50). C(s) represents the concentration (in absorbance units in this case) of species with a given sedimentation coefficient rang.

The densities and viscosities at 25°C were calculated using SEDNTERP for NaCl T-buffer solutions. Partial specific volumes for protein-DNA complexes were calculated using formula: *υ=(υ_DNA_M_DNA_+nυ_n_M_P_)/(M_DNA_+nM_P_)*, where *υ_DNA_* and *M_DNA_* are the partial specific volume and molecular weight of DNA; *υ_P_* and *M_P_* are the partial specific volume and molecular weight of the protein and n is the number of SSB tetramers bound to DNA. In all our experiments the absorbance estimated from integrated c(s) profiles was the same as the absorbance of loaded solutions indicative that no precipitation/aggregation occurred during the experimental runs.

## Results

### Effect of a reversal of backbone ssDNA polarity on *Ec*SSB and *Pf*SSB binding mode formation and cooperativity

Depending on solution conditions one (dT)_70_ is able to bind either one SSB tetramer in its fully wrapped (SSB)_65_ binding mode (65 nts occluded site size) (Fig, 1B) or two SSB tetramers in its (SSB)_35_ binding mode with high cooperativity (Fig, 1C) (20, 24, 51). The proposed structure of an (SSB)_65_ complex in Fig. 1B (6) shows that ~35 nucleotides first wrap around the upper two SSB subunits then DNA crosses the dimer-dimer interface continuing to wrap another 35 nts around the other two subunits following the same path as for the upper subunits. If the SSB tetramer binds ssDNA in a manner that maintains a strict 3’ to 5’ backbone polarity, then a reversal of the backbone polarity in the middle of a (dN)_70_ might weaken or even preclude formation of the fully wrapped (SSB)_65_ binding mode. The ssDNA wrapping topology proposed for the (SSB)_35_ mode, although more speculative (6) (Fig. 1C), also suggests that a reversal of the backbone polarity in the middle of a (dN)_70_ might also affect the relative orientation of two tetramers bound in the (SSB)_35_ mode and affect any cooperative interactions. In order to test these concepts, we designed three non-natural (dT)_70_ variants (see Materials and Methods). In two variants the backbone polarity was reversed in the middle of the DNA by incorporating a 3’-3’ or 5’-5’ phosphodiester linkage; these are denoted as 5’-(dT)_70_-5’ and 3’-(dT)_70_-3’, respectively. The third variant alternated a 3’-3’ and 5’-5’ phosphodiester linkage after each nucleotide and is denoted ap-(dT)_70_. DNA molecules containing backbone polarity reversals have been used to probe directional movement of DNA translocases and helicases (31, 52–57). Binding of these unlabeled DNA molecules was monitored by the quenching of the intrinsic Trp fluorescence of SSB (24, 32, 48). Other versions of these (dT)_70_ variants were made in which the ends of each DNA were labeled with the fluorophores, Cy3 and Cy5, which can undergo Forster resonance energy transfer (FRET). Use of these labeled (dT)_70_ molecules enabled us to monitor formation of the different SSB binding modes since a higher FRET efficiency is observed in the (SSB)_65_ mode due to the closer proximity of the two ends of (dT)_70_ in this mode as demonstrated previously (21, 24).

The results of equilibrium titrations of SSB with (dT)_70_, 5’-(dT)_70_-5’ and 3’-(dT)_70_-3’ monitoring SSB Trp fluorescence quenching are shown in Fig. 2A and 2B in Buffer T at 25.0°C at 10mM and 0.30 M [NaCl]. At 10 mM NaCl, both the (SSB)_65_ and the (SSB)_35_ binding modes can form as determined by the SSB/DNA ratio, whereas at 0.30 M NaCl, only the (SSB)_65_ binding mode is formed (14, 18, 19, 24, 51). At 10 mM NaCl (Fig. 2A) for all (dT)_70_ constructs the normalized quenching increases linearly with addition of DNA indicative of stoichiometric binding (all added DNA binds to SSB). The inflection point at a DNA to protein ratio of 0.5 (r=[SSB]_tot_/[(dT)_70_]_tot_=2) reflects the transition from the (SSB)_35_ mode to the (SSB)_65_ mode, the latter becoming solely populated at r=1 as shown previously for (dT)_70_ (24, 51). However, we note that both RP DNA constructs show a higher extent of Trp quenching in the (SSB)_35_ mode (Q_35_~0.6 for green and orange isotherms in Fig. 2A at [(dT)_70_)]_tot_/[SSB]_tot_=0.5) compared to (dT)_70_ (cyan), for which Q_35_~0.5 is expected (24, 51). This might reflect a slightly different positioning of the two SSB tetramers on the RP DNAs. However, independent of polarity, no difference in quenching (Q_65_=0.9) is observed for the 1:1 (SSB)_65_ complexes that form at [(dT)_70_)]_tot_/[SSB]_tot_>1. At the higher salt concentration of 0.30 M NaCl (Fig. 2B) all three titrations are identical, showing a linear increase in Trp quenching up to r=1 indicative of stoichiometric binding exclusively in the (SSB)_65_ mode (24, 51).

We next performed titrations of the Cy3/Cy5 fluorescently labeled (dT)_70_ constructs with SSB at the same two [NaCl]. The results are consistent with the Trp fluorescence titrations in Fig. 2A and 2B. At 10 mM NaCl (Fig. 2C), the Cy5 FRET signal increases linearly up to r=1 reflecting formation of the fully wrapped (SSB)_65_ complex, and then decreases linearly up to r=2 reflecting a transition to a 2:1 (SSB)_35_ complex with increasing [SSB]. Interestingly, the Cy5 FRET signal is similar for all 1:1 complexes (F_Cy5,65_ ~ 3.5×10^5^ at r=1), whereas the 2:1 complexes (r>2) show different Cy5 FRET signals. SSB bound to 5’-(dT)_70_-5’ shows a higher Cy5 fluorescence compared to 3’-(dT)_70_-3’ and (dT)_70_. This suggests some difference in the DNA wrapping in the (SSB)_35_ mode with 5’-(dT)_70_-5’. However, the sharpness of the transitions between binding modes at r=1 argues that SSB still binds to both RP DNA constructs with a high nearest neighbor cooperativity (24). However, all three titrations at 0.30 M NaCl (Fig. 2D) are identical indicating that they can all form a fully wrapped 1:1 (SSB)_65_ complex in these conditions consistent with the results in Fig. 2B.

To obtain an independent check on our conclusion that both 1:1 and 2:1 SSB complexes can be formed on both RP (dT)_70_ constructs at low [NaCl], we performed sedimentation velocity experiments at total concentration ratios of r=1 and 2 SSB/DNA. Fig. S1A shows the results plotted as c(s) distributions (24, 50). All three DNA molecules sediment identically in the absence of SSB. At each SSB/DNA ratio, only a single c(s) peak is observed (s_20,w_=5.1S at r=1 and s_20,w_=6.7S at r=2). The sedimentation coefficients of the r=1 and r=2 complexes also are the same for each DNA-SSB complex indicating that the change in RP does not lead to major differences in hydrodynamic properties. At the higher 0.30 M [NaCl] (Fig. S1B) only one peak at s_20,w_=5.3S is observed at both r=1 and r=2 indicating that only a 1:1 (SSB)_65_ complex forms at both SSB ratios.

We showed previously that the SSB from *Plasmodium falciparum, Pf*SSB, binds ssDNA only in an (SSB)_65_ binding mode independent of [NaCl] with a DNA topology similar to *Ec*SSB, but with a DNA backbone polarity opposite to that observed in the *E. coli* SSB-DNA structure(41, 45). Using equilibrium titrations (Fig. S2) and sedimentation velocity (Fig. S3) we demonstrate that the backbone polarity of (dT)_70_ has no effect on the formation of the 1:1 fully wrapped complexes, which form stoichiometrically both at low (10 mM) and moderate (0.30 M) NaCl concentrations (s_20,w_=4.8S and s_20,w_=5.4S, respectively, Fig. S3). However, a somewhat lower Cy5 fluorescence (FRET) characterizes formation of the *Pf*SSB-3’-(dT)_70_-3’ complex compared to the 5’-(dT)_70_-5’ complex and (dT)_70_ (Fig. S2). This suggests a slight difference in DNA wrapping in the *Pf*SSB-3’-(dT)_70_-3’ (SSB)_65_ binding mode.

So far we investigated the binding mode formation at NaCl concentrations of salt, where either the (SSB)_35_ ([NaCl]≤0.01 M) or the (SSB)_65_ ([NaCl]≥0.3 M) modes are formed predominantly. However, at intermediate [NaCl] both modes are expected to be populated (14, 15). To monitor the transition between binding modes we performed titrations of DNA constructs with *Ec*SSB at 40, 80 and 100 mM NaCl (Fig. 3) and found that, while the binding of the first SSB tetramer (formation of 1:1 (SSB)_65_ complex) is stoichiometric for all DNAs, the transition to the (SSB)_35_ mode upon binding the second SSB tetramer is less favorable for 5’-(dT)_70_-5’ (green circles), whereas the isotherms for binding 3’-(dT)_70_-3’ and (dT)_70_ are very similar. This difference can be due to either a higher affinity (K_65_) of *Ec*SSB to 5’-(dT)_70_-5’ in the (SSB)_65_ binding mode or a lower nearest-neighbor cooperativity (ω_35_) in the (SSB)_35_ binding mode (24).

**Figure 3.**
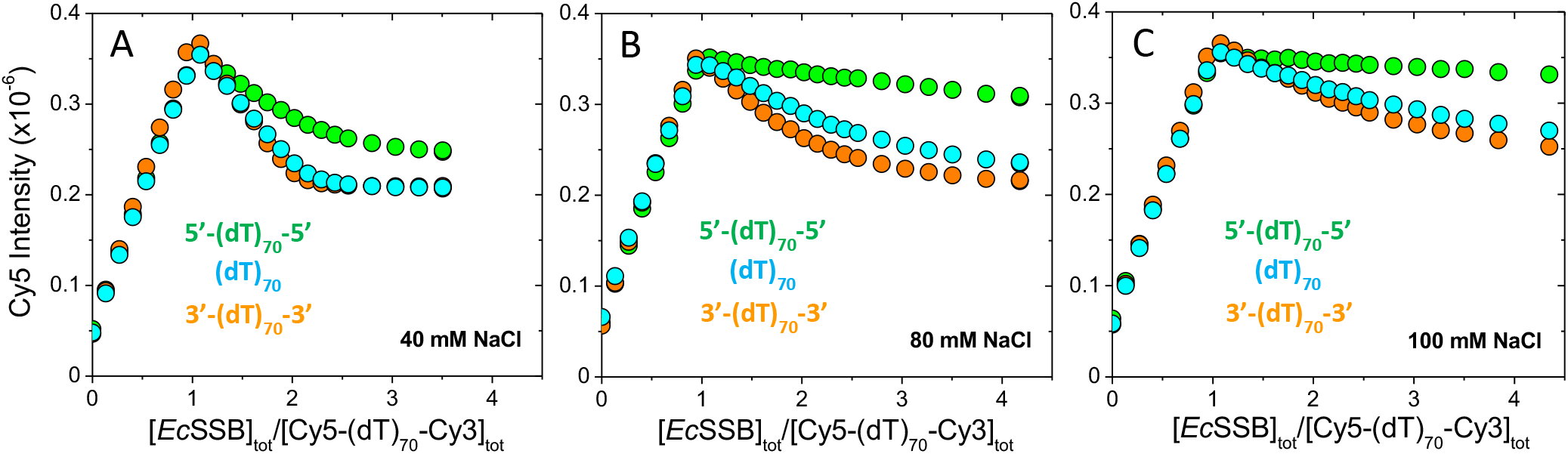
*Ec*SSB binding to reverse polarity (dT)_70_ at intermediate [NaCl]. Results of equilibrium titrations of Cy3/Cy5-labeled (dT)_70_ (cyan), 3’-(dT)_70_-3’ (orange) and 5’-(dT)_70_-5’ (green) (0.1 μM each) with *Ec*SSB monitoring Cy5 fluorescence enhancement (exc: 515 nm, em: 665 nm) in 40 mM NaCl **(A)**, 80 mM NaCl **(B)** and 100 mM NaCl **(C)** (buffer-T, 25°C), plotted as Cy5 fluorescence signal versus the ratio of concentrations of total SSB tetramer to total DNA.

To examine this further, we performed equilibrium titrations of *Ec*SSB with the RP (dT)_70_ DNA constructs (Fig. 4A) in the presence of 2.0 M NaBr, which lowers the binding affinity of *Ec*SSB to (dT)_70_ into a range where it can be measured accurately(34, 51, 58). We find that K_65_ for SSB binding to (dT)_70_ (1.8±0.1)x10^7^ M^-1^) and 3’-(dT)_70_-3’ (3.3±0.1)x10^7^ M^-1^) are similar under these conditions, whereas binding to 5’-(dT)_70_-5’ (2.4±0.1)x10^6^ M^-1^) is much lower. Additional stopped-flow experiments (Fig. S4A) established that the decrease in affinity for 5’-(dT)_70_-5’ is due to a more than 13-fold increase in the dissociation rate constant (k_-5’5’_=13.1±1.9 s^-1^ vs k_-3’3’_=-0.1±0.7 s^-1^, assuming an upper bound value for the latter), whereas the same bimolecular association rate constant of ~ 4×10^7^ M^-1^s^-1^ is measured for both DNA constructs. The lower value of K_65_ for the 5’-(dT)_70_-5’ construct in 2.0 M NaBr suggests that the preference for the (SSB)_65_ binding mode for the 5’-(dT)_70_-5’ construct (see Fig. 2) results from a significantly decreased cooperativity in the (SSB)_35_ mode, (ω_35_).

**Figure 4.**
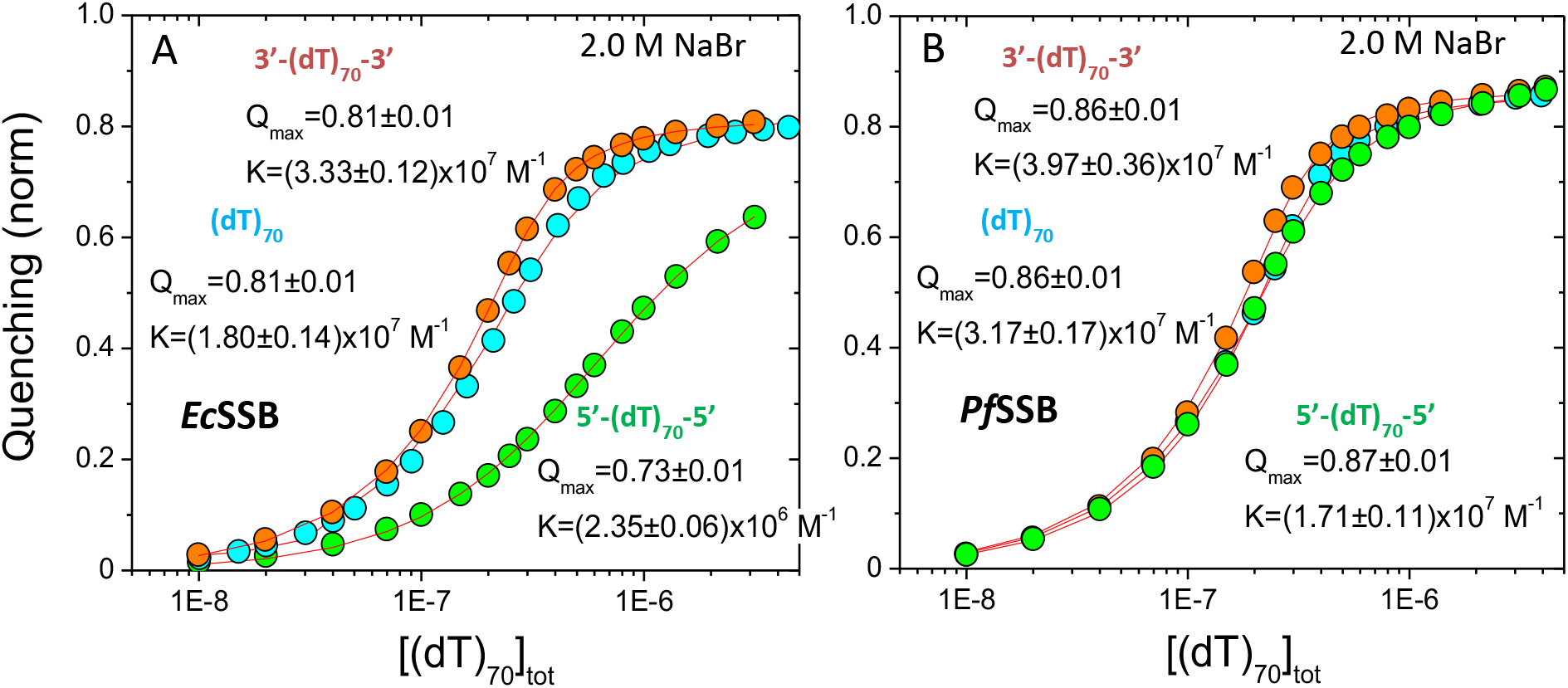
Reverse polarity (dT)_70_ bind to *Ec*SSB with different affinities, but with the same affinity to *Pf*SSB. Results of equilibrium titrations of *Ec*SSB **(A)** and *Pf*SSB **(B)** (0.30 μM each) with (dT)_70_ (cyan), 3’-(dT)_70_-3’ (orange) and 5’-(dT)_70_-5’ (green) monitoring intrinsic Trp fluorescence quenching (ex: 296 nm, em: 350 nm) in 2.0 M NaBr (buffer-T, 25°C). Solid lines are simulated isotherms based on the best fit parameters shown that were obtained from a NLLS fit to a single site binding model (Eqs 1–2 in Materials and Methods).

In contrast, equilibrium binding of *Pf*SSB under the same conditions (see Fig. 4B) shows little difference in affinity for (dT)_70_ (K_65_ =(3.2±0.2)x10^7^ M^-1^) and the two RP DNAs (K_65, 5’5’_=(1.7±0.1)x10^7^ M^-1^ vs. K_65, 3’3’_=(4.0±0.4)x10^7^ M^-1^). Furthermore, *Pf*SSB (Fig. S4B) shows only a ~3 fold difference in dissociation rate constants, k_-5’5’_=3.5±0.5 s^-1^ and k_-3’3’_=1.1±0.7 s^-1^ and similar association rate constants (~3×10^7^ M^-1^ s^-1^). We note that affinities obtained from these rate constants (K=k_on_/k_off_) agree well with the affinities determined from our equilibrium titration experiments (Fig. 4).

In spite of the fact that the affinity in the (SSB)_65_ binding mode might vary for the RP constructs, the most interesting result is that both *Ec*SSB and *Pf*SSB are still able to form fully wrapped complexes with the RP DNA variants under all conditions examined. Furthermore, these RP constructs are also able to form highly cooperative (SSB)_35_ complexes. However, we note that quantification of the nearest neighbor cooperativity parameters for the different DNA constructs is difficult due to the strong correlation of the binding parameters (K_65_, K_35_ and ω_35_). We have recently performed such an analysis for wt *Ec*SSB binding to (dT)_70_, but this required having reasonable estimates of either K_65_ or K_35_(24). Such an analysis cannot be performed at this time since estimates of K_35_ and K_65_ are not available for SSB binding to 5’-(dT)_70_-5’ and 3’-(dT)_70_-3’.

We next performed isothermal titration calorimetry (ITC) experiments in order to probe the thermodynamics of the *Ec*SSB and *Pf*SSB binding to the RP-(dT)_70_ DNA in more detail. Fig. 5 shows the results of titrations of *Ec*SSB (1 μM tetramer) with 5’-(dT)_70_-5’, 3’-(dT)_70_-3’ and (dT)_70_ at low [NaCl] (buffer H, 20 mM NaCl). All *Ec*SSB titrations (Fig. 5A and 5B) show biphasic character, reflecting the initial formation of a 2:1 (SSB)_35_ complex in the range of DNA to protein ratios ≤0.5, which then transforms to a 1:1 (SSB)_65_ complex for DNA to protein ratios from 0.5 – 1.0. Since binding is stoichiometric for each DNA under these conditions, the normalized heats in the flat portions of the isotherms can be related to the enthalpies for formation of the binding modes as given in Eqs. 3 (59),

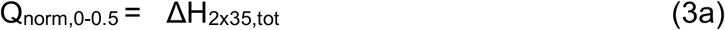

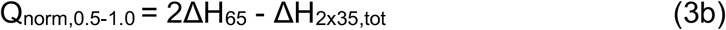

where ΔH_2×35,tot_ =2ΔH_35_ + ΔHω_35_ represents overall enthalpy of (SSB)_35_ binding mode formation (including an enthalpic contribution from cooperativity) and ΔH_65_ is the enthalpy of SSB binding in (SSB)_65_ binding mode. Inspection of Fig. 5A and 5B indicates that *Ec*SSB binding to 3’-(dT)_70_-3’ is very similar to its binding to (dT)_70_ (Fig. 5B) and slightly more enthalpically favorable compared to binding to 5’-(dT)_70_-5’ (Fig. 5A). Solution of the system of equations (Eq. 3) enables us to quantify the differences in the binding enthalpies for the different binding modes formed on the different DNAs. For 5’-(dT)_70_-5’, ΔH_2×35,tot_ = −142.3 ±0.4 kcal/mol and ΔH_65_ = −148.4 ±1.6 kcal/mol, whereas for 3’-(dT)_70_-3’ and (dT)_70_ ΔH_2×35,tot_ = −152.3 ±1.9 kcal/mol and ΔH_65_ = −160.0 ±2.4 kcal/mol, Hence, the interaction of *Ec*SSB with 5’-(dT)_70_-5’ is ~ 10 and 12 kcal/mol less enthalpically favorable in the (SSB)_35_ and (SSB)_65_ binding modes, respectively. This somewhat less favorable binding to 5’-(dT)_70_-5’ is consistent with the equilibrium titrations and stopped-flow results discussed above.

**Figure 5.**
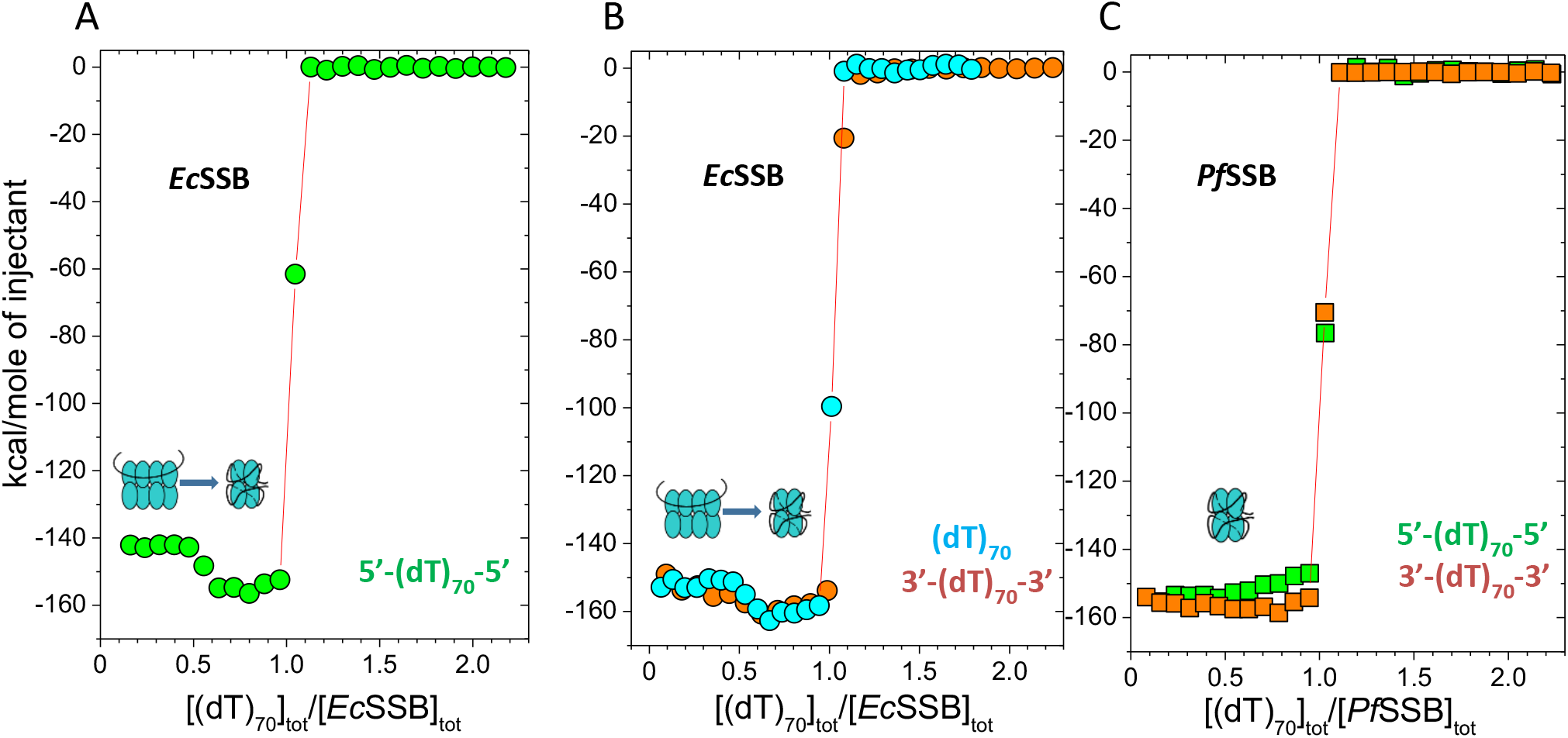
Enthalpic contributions to binding of *Ec*SSB and *Pf*SSB to reverse polarity (dT)_70_. Results of ITC titrations of **(A)-** *Ec*SSB with 5’-(dT)_70_-5’ (green), **(B)-** *Ec*SSB with (dT)_70_ and 3’-(dT)_70_-3’ (cyan and orange), and **(C)-** *Pf*SSB with 3’-(dT)_70_-3’ and 5’-(dT)_70_-5’ (orange and green) in 20 mM NaCl (buffer-H, 25°C) plotted as heats per injection normalized per amount of injected DNA (Q_norm_, kcal/mole) versus the ratio of total DNA to total SSB tetramer concentration. The cartoons depict the (SSB)_35_ and (SSB)_65_ complexes and where they form during the course of the titrations.

In contrast, the interaction of *Pf*SSB with both RP (dT)_70_ constructs at these low salt conditions (Fig. 5C) leads to formation of only the 1:1 (SSB)_65_ complex, consistent with previous studies showing that *Pf*SSB does not form an (SSB)_35_ complex (45). *Pf*SSB binds to DNA molecules with similar enthalpies of ΔH_65,5’5’_ = −152.3 ± 2.3 kcal/mol and ΔH_65,3’3’_ = −156.2 ±1.4 kcal/mol.

### SSB binding to oligodeoxythymidylates with alternating polarity (ap-(dT)_70_ and ap-(dT)_35_)

We next determined whether SSB is able to bind to ssDNA in which the backbone polarity is reversed after each nucleotide. We refer to these as ap-(dT)_70_ (ap-for alternating polarity), ap-(dT)_35_ and Cy5-ap-(dT)_70_-Cy3 DNA (See Materials and Methods). The results are presented in Fig. 6A and 6B, at [NaCl] in the range from 10 mM to 0.30 M. Titrations at 10, 30 and 100 mM NaCl show transition points at DNA to protein ratios of 0.5 and 1.0 indicating that both 2:1 (SSB)_35_ and 1:1 (SSB)_65_ modes can form. Whereas, only a 1:1 (SSB)_65_ complex is formed in 0.30 M NaCl. However, we note that the normalized Trp quenching only reaches values in the range Q_obs_=0.62-0.70 ([DNA]_tot_/[SSB]_tot_>1.0), much lower than the value of Q_65_=0.9 observed for the (SSB)_65_ mode on (dT)_70_ (24, 51) (see also Fig. 2B). This suggests that ap-(dT)_70_ is unable to form a fully wrapped (SSB)_65_ binding mode. This conclusion is also supported by the results of titrations with the C3/Cy5 end-labeled ap-(dT)_70_ shown in Fig, 6B. The 1:1 (SSB)_65_ complexes formed in 0.03 M, 0.1 M and 0.3 M NaCl at r=1 have Cy5 intensities in the range (2.5-2.8)x10^5^, while the Cy5 signal intensity for normal (dT)_70_-SSB complex is ~ 3.5×10^5^ (see Fig. 2C and 2D and Fig. 3).

**Figure 6.**
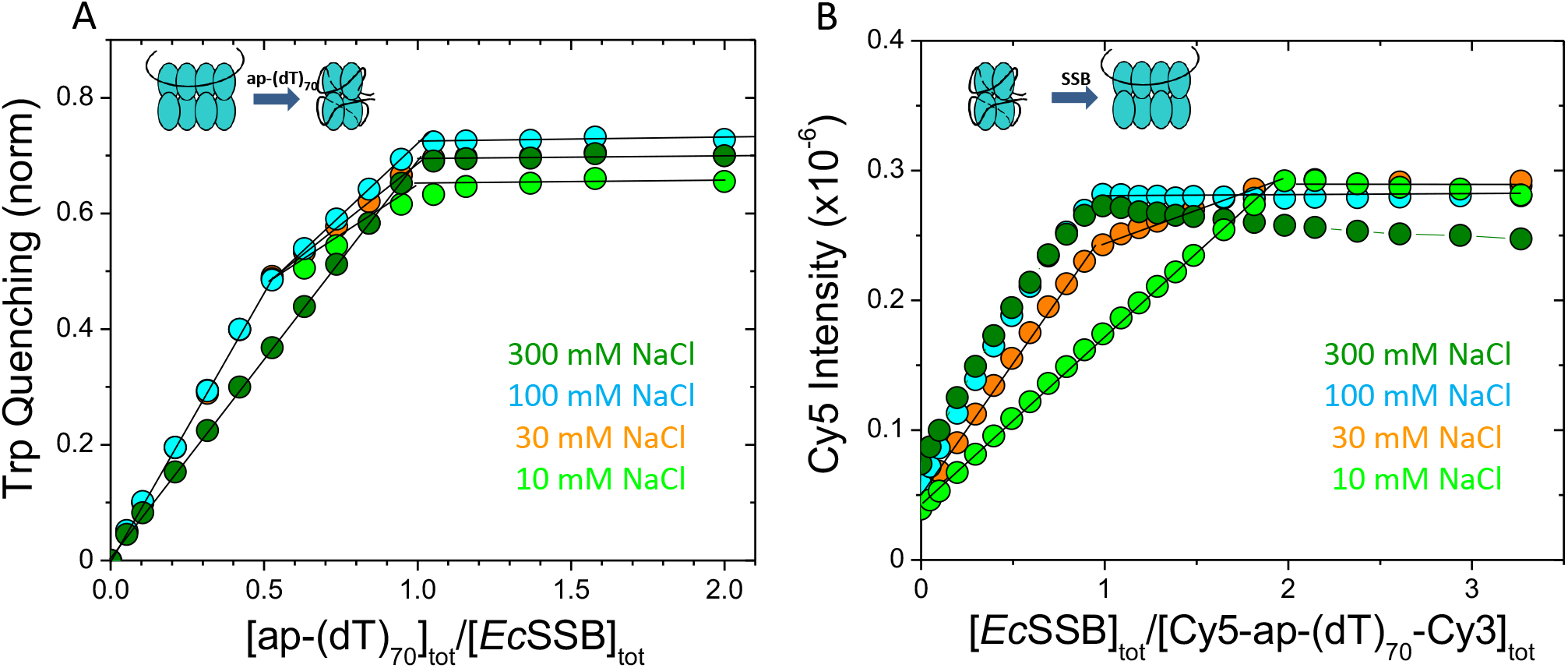
*Ec*SSB binding to alternating polarity (ap)-(dT)_70_. **(A)** - results of equilibrium titrations of *Ec*SSB (0.30 μM) with ap-(dT)_70_ in buffer-T (25°C) in 10 mM NaCl (green), 30 mM NaCl (orange), 100 mM NaCl (cyan) and 300 mM NaCl (dark green), plotted as fractional Trp fluorescence quenching versus the ratio of total DNA to SSB tetramer concentrations. The cartoons depict the (SSB)_35_ and (SSB)_65_ modes and expected transitions during the titrations. **(B)** - results of equilibrium titrations of Cy3/Cy5 labeled ap-(dT)_70_ (0.1 μM each) with *Ec*SSB in 10 mM NaCl (green), 30 mM NaCl (orange), 100 mM NaCl (cyan) and 300 mM NaCl (dark green) in buffer-T (25°C), plotted as Cy5 fluorescence versus the ratio of concentrations of total SSB tetramer to total DNA. The cartoons depict the (SSB)_35_ and (SSB)_65_ modes and expected transitions during the titrations.

The ap-(dT)_70_ data are also consistent with the results for *Ec*SSB binding to ap-(dT)_35_. Two molecules of normal polarity (dT)_35_ can bind to an SSB tetramer, although with negative cooperativity that depends on salt concentration and type (51, 60). All titrations of SSB with ap-(dT)_35_ presented in Fig. S5 show that only one ap-(dT)_35_ molecule binds to an SSB tetramer at all [NaCl]. Binding is stoichiometric in the 0.01-0.1 M NaCl concentration range (panel A) and affinity is measurable (k_1,35_=1.7±0.2)×10^8^ M^-1^) at 0.3 M NaCl (panel B), whereas binding of the first molecule of normal (dT)_35_ is always stoichiometric (k_1,35_ affinity is too high to measure) even at much higher [NaCl] (51, 58). The results shown in Fig. S5B also indicate that even at a ~16 fold molar excess of ap-(dT)_35_ only a small fraction of a 2:1 complex can form (k_2,35_=6.8±0.6)×10^4^ M^-1^ is very low). This inability to bind a second molecule of ap-(dT)_35_ is consistent with inability of ap-(dT)_70_ to form a fully wrapped complex. However, it is surprising that *Ec*SSB still can bind the ap-(dT)_35_ with substantial affinity.

## Discussion

The two tetrameric SSB proteins from *E. coli* and *P. falciparum* have similar subunit organization: an N-terminal DNA binding core and an intrinsically disordered C-terminal tail (Fig. 1A). While the tetrameric DNA binding cores share 39% amino acid identity, and 66% homology (41), their intrinsically disordered C-terminal domains differ greatly. In *Ec*SSB the conserved acidic tip (9 aa) of the C-terminal tail, which serves as a binding site for numerous metabolic SIPs (SSB interaction proteins) (5, 8, 13), is connected to the DNA binding core via a 56 aa linker containing few charged residues (Fig. 1A). In contrast, *Pf*SSB possesses a different acidic tip that lacks the conserved IPF residues of the *Ec*SSB tip, and is connected to the DNA binding core by a longer (82 aa) more highly charged linker. This difference in the C-terminal tails has a dramatic effect on the DNA binding properties, including binding mode formation and cooperative binding (23, 24, 32). In particular the *Pf*SSB is unable to form a low site size, (SSB)_35_ equivalent binding mode (45) and shows no evidence of highly cooperative binding to ssDNA (32) in spite of the fact that the DNA binding cores of *Ec*SSB and *Pf*SSB are very similar.

Our focus in this study was whether *Ec*SSB and *Pf*SSB proteins bind ssDNA with strict polarity. This was motivated by the observation that crystallographic structures show that *Pf*SSB and *Ec*SSB bind ssDNA with opposite polarity and slightly different topology in their fully wrapped (SSB)_65_ binding modes (compare Fig. 1B and 1D). In fact, of the SSB-ssDNA crystal structures that have been reported, only the *Ec*SSB OB-fold binds ssDNA with a different polarity.

To examine this we designed two types of (dT)_70_ variants with modified phosphodiester backbone polarities. In the first variant, a reversal of the backbone polarity, was introduced in the middle of the sequence, either via a 3’-3’ or 5’-5’ linkage. In a second variant, every phosphodiester backbone in the sequence was reversed, i.e., a 3’-3’ linkage alternated with a 5’-5’ linkage, designated ap-(dT)_70_ and ap-(dT)_35_. If either SSB binds ssDNA with strict polarity, we hypothesized that a fully wrapped (SSB)_65_ binding mode would be inhibited. Surprisingly, we found that both 3’-(dT)_70_-3’ and 5’-(dT)_70_-5’ bind the same as normal (dT)_70_ to *Pf*SSB. Only 1:1 fully wrapped complexes with the same hydrodynamic characteristics are formed stoichiometrically at both high (0.30 M) and low (10 mM) [NaCl] (Fig. S2 and S3). Moreover, no differences are observed in the thermodynamics and kinetics of complex formation (Figs. 4B, 5C and S4B).

However, the binding of *Ec*SSB to these variants shows more differences. Both (SSB)_35_ and (SSB)_65_ complexes can form stoichiometrically on the RP (dT)_70_ similar to normal (dT)_70_ at low (10 mM) and high (0.30 M) [NaCl] (Fig. 2 and Fig. S1). However, the relative stabilities of the two modes are affected. At intermediate [NaCl] (40-100 mM) the (SSB)_65_ mode is favored with the 5’-(dT)_70_-5’ RP construct, whereas (dT)_70_ and 3’-(dT)_70_-3’ show little difference (Fig. 3). This could result from increased affinity (K_65_) in the (SSB)_65_ mode or decreased affinity (K_35_) or cooperativity (ω_35_) in the (SSB)_35_ binding mode (24). However, equilibrium titrations (Fig. 4A) and kinetic data (Fig. S4A) in buffer containing 2.0 M NaBr indicate that actually the affinity in the (SSB)_65_ mode (K_65_) is ~10-fold lower for the 5’-(dT)_70_-5’ construct compared with (dT)_70_ and 3’-(dT)_70_-3’. This suggests that the differences are due to weakening of the interactions (either K_35_ and/or ω_35_) of SSB with 5’-(dT)_70_-5’ in the (SSB)_35_ binding mode. This is corroborated by ITC measurements (Fig. 5A and 5B) demonstrating that the total enthalpies for forming both binding modes on 5’-(dT)_70_-5’ are 10-12 kcal/mol less favorable compared with (dT)_70_ and 3’-(dT)_70_-3’. However, even for *Ec*SSB, the effects of the backbone polarity reversal on DNA binding is relatively small.

Why would a backbone polarity reversal in the middle of (dT)_70_ have no effect on formation of the *Pf*SSB (SSB)_65_ binding mode, and show only small effects for *Ec*SSB and only with 5’-(dT)_70_-5’. We hypothesize that this is related to some structural differences of the DNA binding domains (tetrameric cores) of these proteins, which dictate slightly altered pathways of ssDNA wrapping in the two (SSB)_65_ binding modes. In *Ec*SSB the ssDNA wraps around the upper AA’ dimer and then crosses the dimer-dimer interface towards the β helix of the B subunit, finally wrapping around the B’B dimer in *“trans’’* (AA’BB’ topology, Fig. 1B). Whereas in *Pf*SSB, as well as in *HpSSB* (Fig. 1D and 1E, respectively) ssDNA wraps around the upper and lower dimers similarly to *Ec*SSB (although with opposite polarity), but crosses the dimer-dimer interface under the β helix of the A’ subunit in “*cis*” following an different AA’B’B topology. The *Ec*SSB “*trans*’ topology appears to be dictated by the presence of 5-6 rigid loops (6, 61)(Fig. 1B), whereas a different orientation of these loops in *Pf*SSB (41) (Fig. 1D) and their absence in *Hp*SSB (42) (Fig. 1E) promotes wrapping of ssDNA in “*cis*”. These different crossings of DNA at the dimer-dimer interface could explain the differences that we observe for *Ec*SSB and *Pf*SSB. As a matter of fact for the “*cis*” wrapping pathway that is displayed in the *Pf*SSB structure (Fig. 7Ai) could accommodate fully wrapped 3’-(dT)_70_-3’ and 5’-(dT)_70_-5’ constructs in the (SSB)_65_ binding mode, with no change in the polarity of ssDNA binding to the individual OB-folds, as shown in Fig. 7Aii and Fig. 7Aiii, respectively.

**Figure 7.**
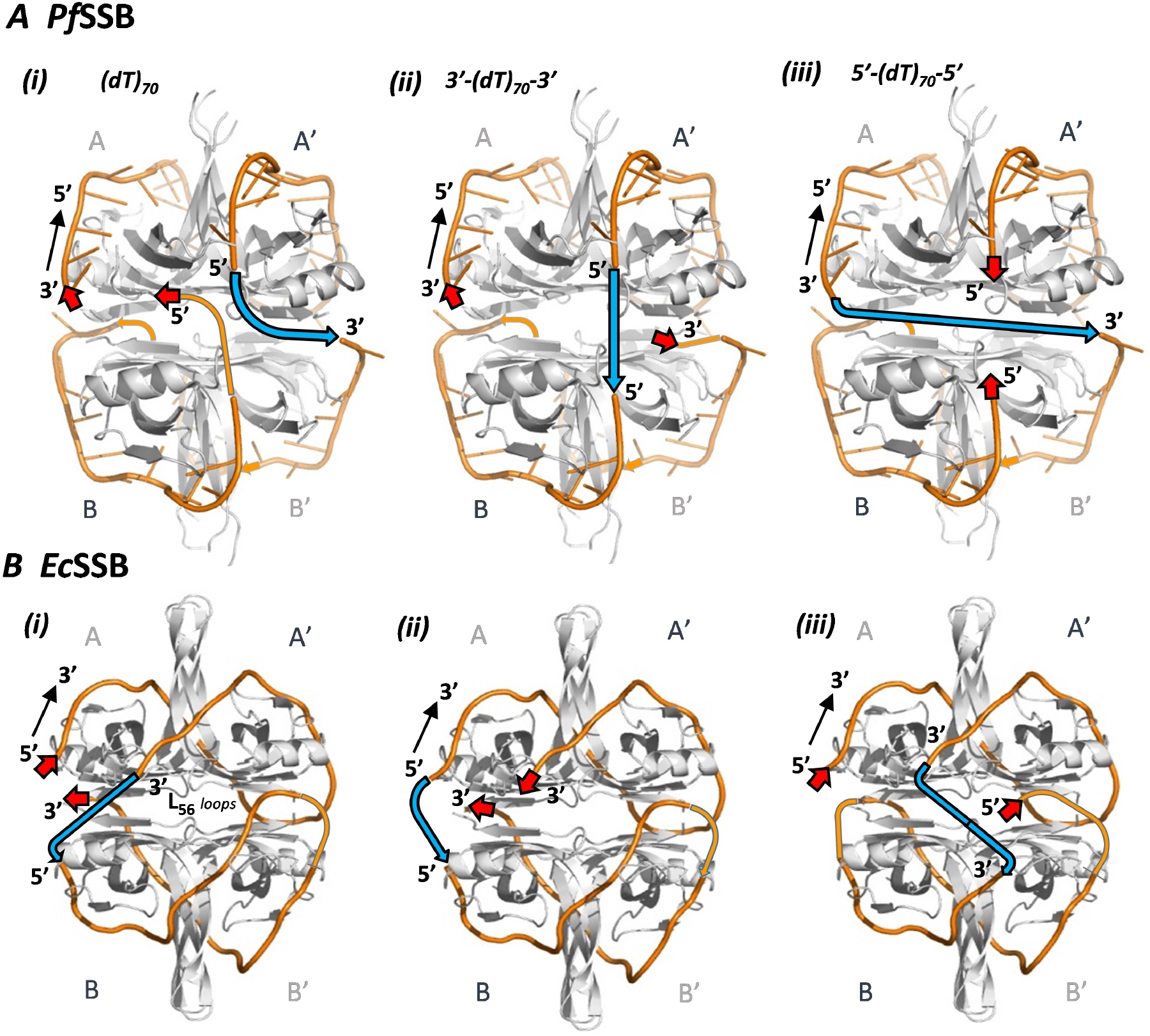
Proposed alternative DNA wrapping topologies in the (SSB)_65_ binding modes for *Ec*SSB and *Pf*SSB formed with the RP (dT)_70_. Proposed topology of ssDNA binding to *Pf*SSB **(A)** and *Ec*SSB **(B)** in an (SSB)_65_ binding mode for (dT)_70_ with normal polarity **(i)**, 3’-(dT)_70_-3’ **(ii)** and 5’-(dT)_70_-5’ **(iii)**. In each case the ends of the DNA entering and exiting the tetramer are designated by red arrows with the indicated polarity. The blue arrows crossing the dimer-dimer (AA’/BB’) interface in each particular case represent the middle regions of the DNA showing the different wrapping topologies needed to maintain the polarity of the DNA backbone bound to each individual subunit (OB-fold) (3’-5’ for *Pf*SSB and 5’-3’ for *Ec*SSB).

The situation differs for *Ec*SSB where DNA follows the “trans” topology (Fig. 7Bi). In this case the DNA strand crossing the dimer-dimer interface (in blue) is further removed from the DNA strand wrapping around the opposite B’ subunit. Assuming that binding polarity to the individual OB-folds is maintained, we propose the wrapping topologies for the 3’-(dT)_70_-3’ and 5’-(dT)_70_-5’ DNA constructs shown in Fig. 7Bii and Fig. 7Biii, respectively. Interestingly, in the proposed topology for the 3’-(dT)_70_-3’ DNA (Fig. 7Bii), the DNA crosses the dimer-dimer interface in the same region as proposed for normal polarity (dT)_70_. However, in the proposed topology for the 5’-(dT)_70_-5’ DNA (Fig. 7Biii), the dimer-dimer crossing occurs in the region of the 5-6 loops, which would appear to be sterically unfavorable. This difference might explain the lower affinity and faster dissociation of 5’-(dT)_70_-5’ from *Ec*SSB, whereas 3’-(dT)_70_-3’ and (dT)_70_ show similar binding properties.

The ability of *Ec*SSB to form a 2:1 (SSB)_35_ complex on both (dT)_70_ RP variants is not as surprising since in that binding mode only two subunits of each bound tetramer interact with the DNA. However, due to different orientation of those tetramers on the RP constructs, a decrease in the nearest-neighbor cooperativity might be expected due to the loss of some contacts required for such interactions. Yet, we still observe significant cooperativity for the 3’-(dT)_70_-3’ DNA, which is comparable to (dT)_70_, whereas cooperativity is decreased with the 5’-(dT)_70_-5’ DNA. Based on a proposed model for the (SSB)_35_ binding mode presented in Fig. 1C, a 2:1 complex should be possible with both RP (dT)_70_ constructs. A simple 180° rotation of the upper tetramer within the x-y plane relative to the lower tetramer creates a 5’-(dT)_70_-5’ complex, while a similar rotation performed on the lower tetramer creates a 3’-(dT)_70_-3’ complex. However, in both cases the interactions between the 4-5 loops that have been proposed to be important for nearest neighbor cooperativity (6) would be lost. We speculate that alternative orientations of the two tetramers in 3D space (e.g. around the Y axes) could restore such nearest-neighbor cooperative contacts and this might be easier for the 3’-(dT)_70_-3’ construct. We note here, that the model for the (SSB)_35_ binding mode in Fig. 1C is only speculative and awaits experimental testing. So alternative wrapping pathways in this mode, which might explain the differences observed for the RP (dT)_70_ constructs cannot be excluded. We also know that nearest neighbor cooperativity in the (SSB)_35_ mode is also dependent on the intrinsically disordered linkers (IDL) in the C-terminal tails (24) that are not observed in the crystal structures. If cooperativity requires direct interactions between the IDLs of adjacent tetramers, such interactions should be possible on (dT)_70_ and both RP variants.

Interestingly, we find that *Ec*SSB still binds stoichiometrically to the fully alternating ap-(dT)_70_ in both binding modes (Fig. 6) over a broad NaCl concentration range. However, the affinity in a fully wrapped (SSB)_65_ mode is weakened significantly. This is indicated by the fact that *Ec*SSB is not able to bind more than one molecule of ap-(dT)_35_ under any solution condition (Fig. S5). Yet, it is somewhat surprising that it can still bind to *Ec*SSB with substantial affinity. E.g. ap-(dT)_35_ binds to *Ec*SSB in 0.30 M NaCl with K_35,1_=1.7×10^8^ M^-1^(Fig. S5B), whereas a lower limit for normal (dT)_35_ is ~ 2×10^9^ M^-1^(51).

The results reported here demonstrate that both tetrameric *Ec*SSB and *Pf*SSB can accommodate stretches of ssDNA with reverse polarity and still bind with high affinity. *Ec*SSB also retains the ability to form its different binding modes and even retains high nearest neighbor cooperativity. Only severe backbone polarity changes, such as for the ap-ssDNA constructs, show a large effect on affinity. We have proposed structural models that can explain this behavior while still retaining strict backbone polarity for binding to the individual OB-folds. Hence, our results are consistent with the different ssDNA polarities observed in the crystal structures of ssDNA complexes with *Ec*SSB and *Pf*SSB. This adaptability may play important roles in the metabolic functions of SSB.

## Author Contributions

TML and AGK designed the research; AGK performed and analyzed the experiments; AGK and TML wrote the manuscript

## Acknowledgments

We thank Roberto Galletto for discussions and comments on the ms. and T. Ho for synthesis and purification of oligodeoxynucleotides. This research was supported in part by the NIH (R01GM30498 and R35GM136632 to TML).

