## Supplementary Material for "Probing *E. coli* SSB Protein-DNA topology by reversing DNA backbone polarity"

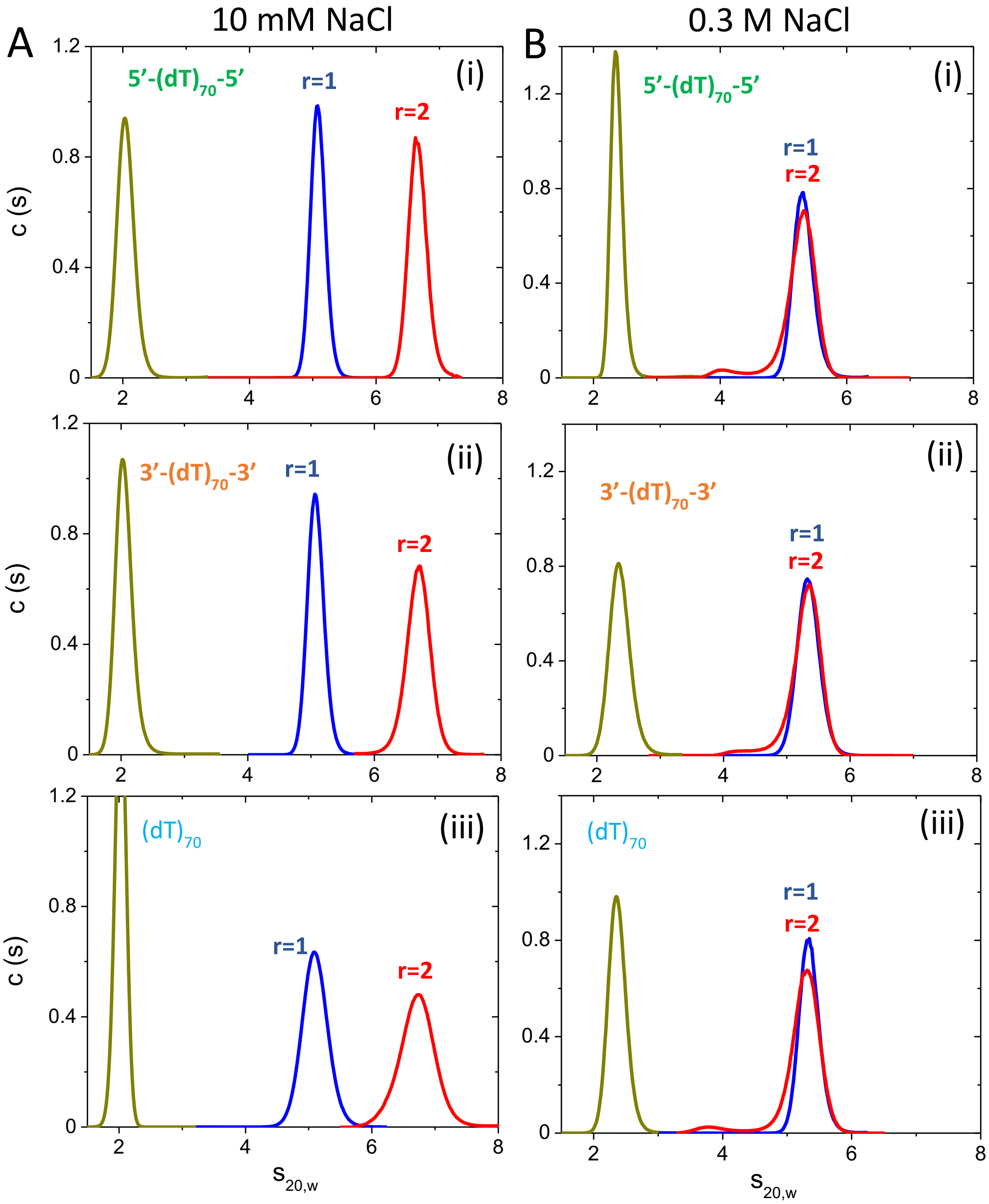

**Figure S1. Sedimentation velocity data for *Ec*SSB support formation of $(SSB)_{35}$ and $(SSB)_{65}$ binding modes on reverse polarity $(dT)_{70}$ constructs similar to normal polarity DNA.**

c(s) profiles from sedimentation velocity experiments performed in the presence 10 mM NaCl **(A)** and 300 mM NaCl **(B)** (buffer-T, 25˚C) for *Ec*SSB complexes with 5′-$(dT)_{70}$-5′ **(i)**, 3′-$(dT)_{70}$-3′ **(ii)**, and $(dT)_{70}$ **(iii)**, formed at different protein to DNA ratios, $r=[SSB]_{tot}/[dT_{70}]_{tot}$ ($[dT_{70}]_{tot}$=0.5 μM): r=0, DNA alone (dark yellow); r=1.0 (blue) and r=2.0 (red). The profiles are converted to 20˚C, water conditions as described in Materials and Methods and reflect formation of both 1:1 $(SSB)_{65}$ ($s_{20,w, AVER}$ = 5.09 ± 0.02 S) and 2:1 $(SSB)_{35}$ ($s_{20,w, AVER}$ = 6.68 ± 0.02 S) complexes in 10 mM NaCl **(A)** and the only one 1:1 $(SSB)_{65}$ complex ($s_{20,w, AVER}$ = 5.31 ± 0.03 S) in 300 mM NaCl **(B)**. The error was estimated as SD.

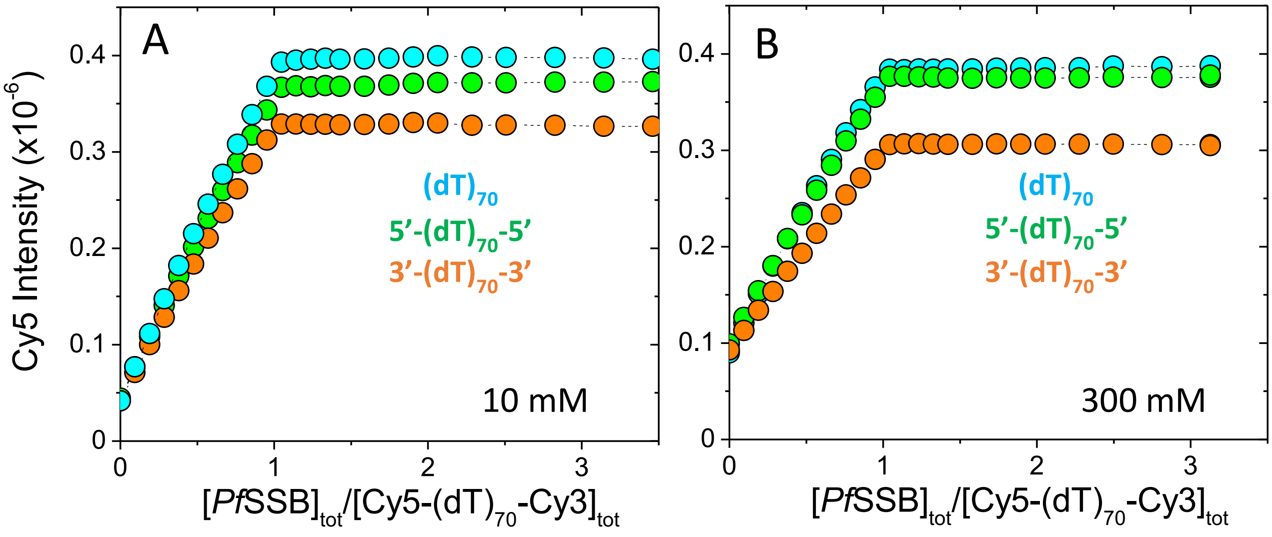


**Figure S2. Equilibrium titration data indicate that *Pf*SSB forms only 1:1 complex with $(dT)_{70}$ reverse polarity constructs independent of [NaCl].**

Results of equilibrium titrations of $(dT)_{70}$ (cyan), 3′-$(dT)_{70}$-3′ (orange) and 5′-$(dT)_{70}$-5′ (green) labeled with Cy3/Cy5 FRET pair (0.1 μM each) with *Pf*SSB monitoring Cy5 fluorescence enhancement (exc: 515 nm, em: 665 nm) in 10 mM NaCl (A) and 0.30 M NaCl (B) (buffer-T, 25$^{o}$C) plotted as Cy5 fluorescence signal at each point of the titration versus the ratio of concentrations of total SSB tetramer to total DNA.

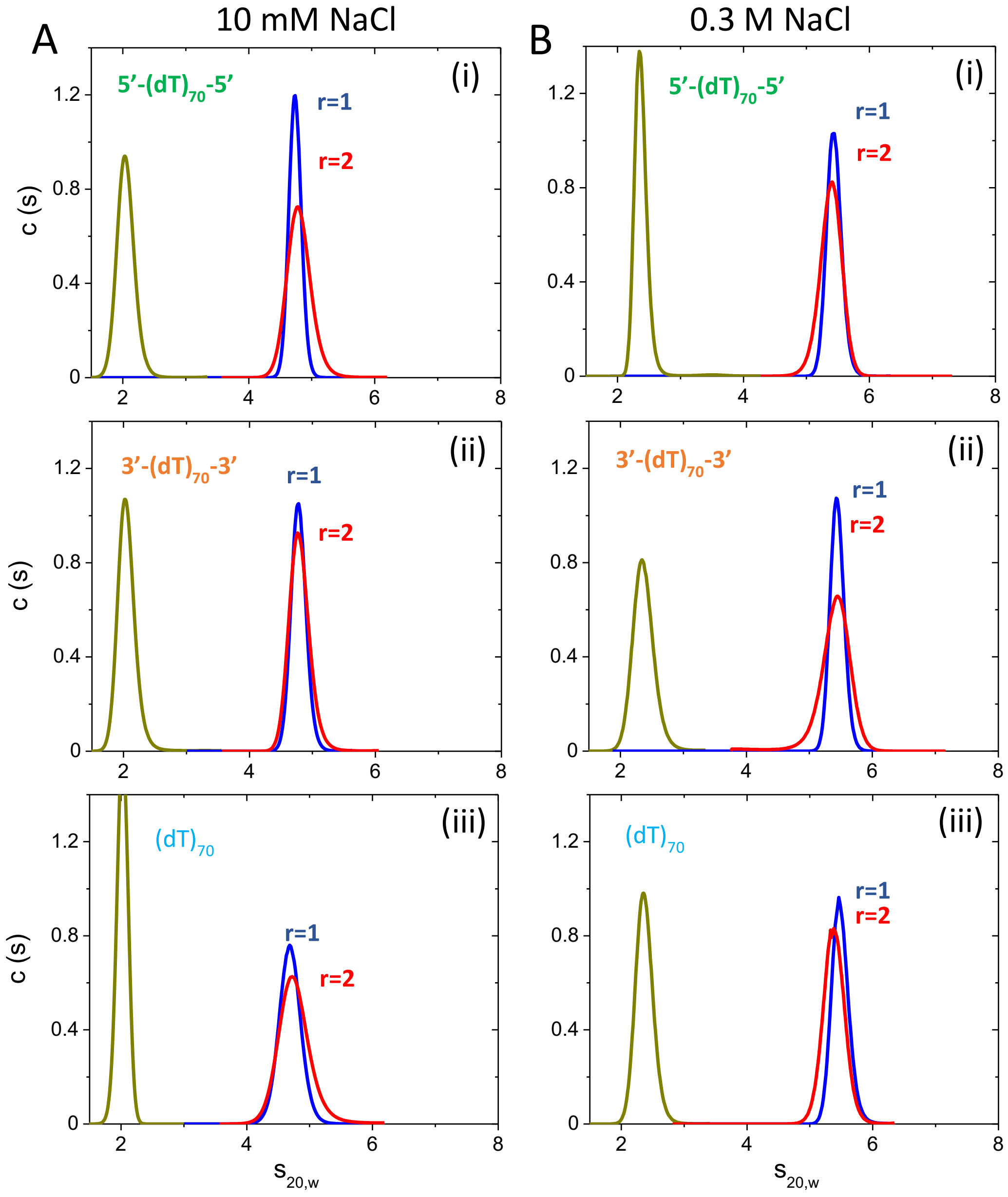


**Figure S3. Sedimentation velocity data for *Pf*SSB support formation of only 1:1 complex on all $(dT)_{70}$ constructs independent of [NaCl].**

C(s) profiles from sedimentation velocity experiments performed in the presence 10 mM NaCl **(A)** and 300 mM NaCl **(B)** (buffer-T, 25ºC) for *Pf*SSB complexes with 5′-$(dT)_{70}$-5′ **(i)**, 3′-$(dT)_{70}$-3′ **(ii)**, and $(dT)_{70}$ **(iii)**, formed at different protein to DNA ratios, $r=[SSB]_{tot}/[(dT)_{70}]_{tot}$ ($[(dT)_{70}]_{tot}$=0.5 μM): r=0, DNA alone (dark yellow); r=1.0 (blue) and r=2.0 (red). The profiles are converted to 20°C, water conditions as described in Materials and Methods and reflect formation of the only one 1:1 fully wrapped complex at both salt concentrations ($s_{20,w,\,AVER}$ =4.76 ± 0.04 S in 10 mM NaCl and $s_{20,w,\,AVER}$ = 5.42 ± 0.03 S in 300 mM NaCl). The error was estimated as SD.

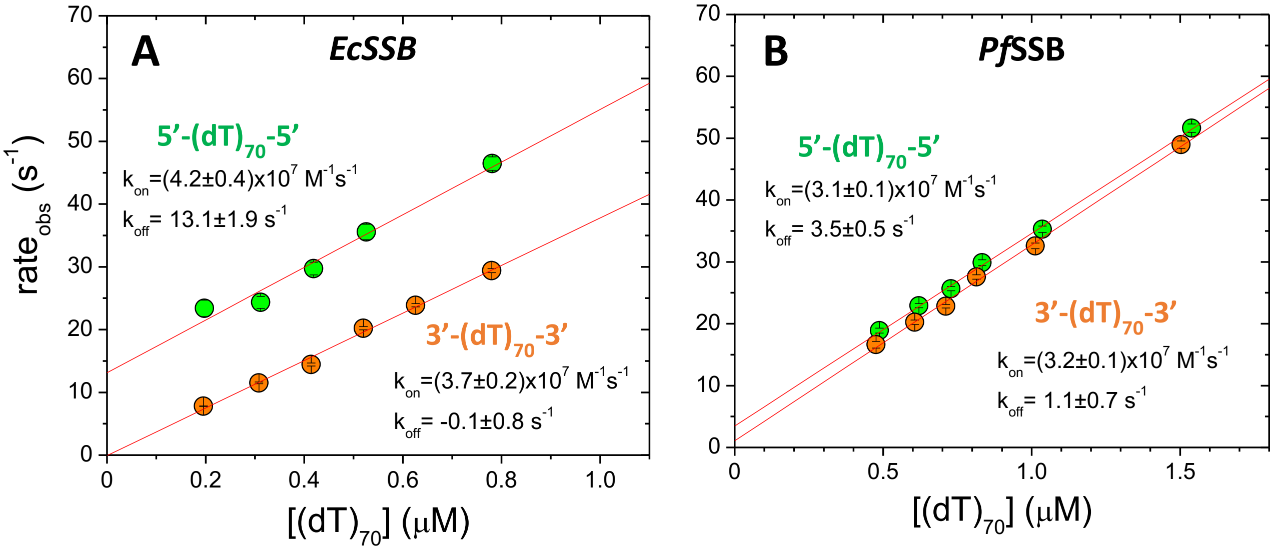


**Figure S4. Stopped-flow experiments suggest that *Ec*SSB dissociates faster from its 1:1 complex with 5'-(dT)$_{70}$-5', whereas *Pf*SSB kinetics shows little difference between RP constructs.**

The dependencies of observed rates ($r_{obs}$) on the concentrations of 5′-(dT)$_{70}$-5′ (green) and 3′-(dT)$_{70}$-3′ (orange) (buffer-H, 2.0 M NaBr, 25°C) obtained adding increasing amounts of DNA to 50 nM of *Ec*SSB **(A)** and *Pf*SSB **(B)** and fitting time courses of Trp fluorescence quenching (exc: 295 nm, em: 335 nm long pass filter) to single exponential decay function for each point of titration. Linear fits of these dependencies produced values of $k_{on}$ and $k_{off}$ (as slope and intercept, respectively) which are presented in corresponding figure panels.

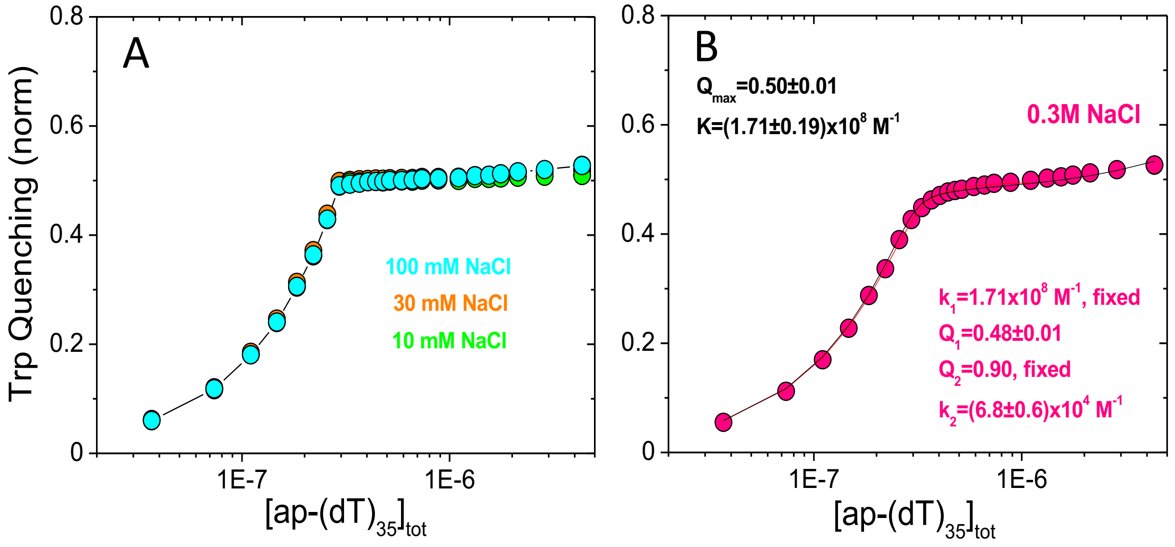


**Figure S5. *Ec*SSB is able to bind only one molecule of ap-$\underline{(dT)_{35}}$ suggesting that $(SSB)_{65}$ binding mode is difficult to form on the substrates with fully alternating polarity.**

Results of equilibrium titrations of *Ec*SSB (0.30 µM) with ap-$(dT)_{35}$ monitoring intrinsic Trp fluorescence quenching (ex: 296 nm, em: 350 nm) in buffer-T (25°C) in the presence of 10 mM NaCl (green), 30 mM NaCl (orange), 100 mM NaCl (cyan) **(A);** and 300 mM NaCl **(B)**, plotted as normalized Trp fluorescence quenching $Q=(F_0\text{-}F_{obs})/F_0$ versus the ratio of total DNA to SSB tetramer concentrations (where $F_0$ is the fluorescence intensity of wtSSB alone and $F_{obs}$ is the observed Trp fluorescence measured at each point in the titration and $Q_{max}=(F_0\text{-}F_{max})/F_0)$, where $F_{max}$ is fluorescence at saturation). At [NaCl]≤0.1 M **(A)** only one $(dT)_{35}$ molecule stoichiometrically binds to *Ec*SSB (the affinity is too high to be determined), whereas at 0.3 M NaCl **(B)** the affinity is weaker for the binding of the first molecule and the isotherm shows indication of the second molecule binding. For this case, the fit of the first 19 points of the isotherm to one binding site model, described in Materials and Methods (thin red line) provides the following estimates of the parameters for 1:1 complex formation: $K=(1.71\pm0.19)\times10^{8}$ $M^{-1}$ and $Q_{max}=0.50\pm0.0$. The estimate of the affinity for much weaker binding of the second $(dT)_{35}$ ($k_{2,35}=(6.8\pm0.6)\times10^{4}$ $M^{-1}$) is obtained fitting the data to 2 sequential sites binding model described below (thin black line), when fixing $k_{1,35}=1.71\times10^{8}$ $M^{-1}$ and $Q_2=0.9$ and floating $k_{2,35}$ and $Q_1$, where $Q_2=0.9$ is the characteristic value of normalized Trp fluorescence quenching expected for the *Ec*SSB with two $(dT)_{35}$ molecules bound (32, 51, 48). The errors were estimated as SD.

**Data analysis.**

Binding isotherms in Fig. S5B were analyzed using a single site binding model (see Materials and Methods) and two-site sequential binding model described in Eq. 1a,

$$Q = \frac{Q_1 2k_1 X + Q_2 kk_2 X^2}{1 + 2k_1 X + k_1 k_2 X^2} \qquad (1a)$$

where $Q$ is the observed normalized fluorescence quenching and $Q_1$ and $Q_2$ are the fluorescence quenching corresponding to one and two $(dT)_{35}$ bound, respectively; $k_1$ and $k_2$ are the observed step-wise microscopic association constants for the binding of the first and the second DNA molecule. The concentration of free DNA, $X$, was determined from mass conservation Eq. 1b,

$$X_{tot} = X + X_{bound} = X + \frac{2k_1 X + 2k_1 k_2 X^2}{1 + 2k_1 X + k_1 k_2 X^2} M_{tot} \qquad (1b)$$

where $X_{tot}$ and $M_{tot}$ are total concentrations of $(dT)_{35}$ and protein, respectively. NNLS fitting of the isotherms to Eqs. 1 to obtain the binding parameters was performed using SCIENTIST (Micromath, St Louis, MO) as described (48).
